# NanoLabel: A fast and accurate real-time nanopore signal classifier

**DOI:** 10.64898/2026.05.03.722500

**Authors:** Daanish Mahajan, Chirag Jain, Navin Kashyap

**Affiliations:** Interdisciplinary Mathematics Initiative, Indian Institute of Science, CV Raman Rd, 560012, Karnataka, India; Department of Computational and Data Sciences, Indian Institute of Science, CV Raman Rd, 560012, Karnataka, India; Department of Electrical Communication Engineering, Indian Institute of Science, CV Raman Rd, 560012, Karnataka, India

**Keywords:** Adaptive sampling, enrichment, supervised classification

## Abstract

Oxford Nanopore Technologies’ adaptive sampling capability promises to reduce sequencing cost and turnaround time. At its core, adaptive sampling is a real-time classification problem that distinguishes reads originating from regions of interest. Direct signal-based classification approaches bypass the computational bottleneck of basecalling and can eliminate the need for powerful GPUs. However, operating directly on noisy raw signals remains challenging in real-time settings, where classification decisions must be made quickly. In this work, we propose NanoLabel, a new method for real-time classification of nanopore signals. We build NanoLabel on top of signal-based read mapping tool, RawHash2. We accelerate the classification workflow by mapping reads using only the target regions as the reference. To further improve accuracy, we train a lightweight classifier on mapping-derived features and introduce a data augmentation strategy to construct sufficiently large and class-balanced training datasets. We evaluate NanoLabel using publicly available real sequencing datasets from three human genomes (HG001, HG002, and HG005), while assuming a cancer gene panel as the target. Compared to directly mapping reads with RawHash2, we demonstrate 80 × improvement in the classification time and 0.10 *-* 0.25 units improvement in the F1 score.

## 1. Introduction

Long-read whole-genome sequencing using Oxford Nanopore Technology (ONT) produces reads of average length greater than 10 kbp. However, many biological studies focus only on selected genomic loci, such as detecting pathogenic sequences in samples dominated by host DNA [18] or specific genes within a genome [20]. In such settings, sequencing non-target regions leads to unnecessary sequencing costs and increased computational burden during downstream analyses.

Biochemical techniques, such as PCR [26], hybrid capture [4], and CRISPR/Cas9 [12], have been employed for target enrichment; however, these methods require more time and expertise. Also, this process of amplification erases the information about nucleotide modifications [21, 26]. ONT provides an automated method for achieving targeted sequencing. The DNA bases passing through the nanopore affect the flow of ionic current in the central constriction of the pore, which is recorded by the device. The changes in electrical current depend on the specific chemical properties of the nucleotides, including secondary structure interactions and epigenomic modifications such as methylation. Each channel can individually reverse the voltage across its pore, rejecting the molecule if unnecessary. This creates more room for the target molecules to get sequenced. One can request enrichment of the target or depletion of unwanted DNA molecules directly during sequencing by simply providing a target DNA reference sequence [18].

The process of accepting or rejecting is controlled by the Read-Until API. The signal is analyzed in a streaming fashion by examining small, consecutive, non-overlapping windows called *chunks*. A chunk typically consists of about 4000 current measurements, which translates to approximately 420 bp after basecalling. The API attempts to make the decision with as few chunks as possible. After the first chunk is read, it is basecalled to obtain a nucleotide sequence and then mapped to the reference genome [20]. If the mapper is not confident about the location of origin of the data, the next chunk is processed in a similar fashion, and the information from all the previous chunks is combined to make the decision again. This process continues until the confidence score of mapping exceeds a certain threshold or the read length is insufficient, in which case the read remains unmapped.

To achieve sufficient enrichment, the decision to accept or reject needs to be made quickly because of the speed at which the read passes through the pore, which is approximately 400 nt/sec with R10.4.1 pore chemistry. A recent study [6] showed that basecalling is a bottleneck in the Read-Until pipeline. Also, basecallers are computationally intensive and require powerful GPUs for operation [3]. This limits the application of classification on basecalled data in scenarios with constrained computational resources. Hence, demanding fast, accurate, basecalling-free adaptive sampling solutions. Deep learning models have shown promising results on inter-species classification tasks [2, 23, 17], but fail to distinguish well on intra-species classification tasks, especially for the human genome [23]. Consequently, alignment-based techniques that explicitly account for the repetitive structure of the human genome represent a more promising alternative.

One way to address the classification problem is to find an approximate nucleotide representation of the input signal and use it for mapping. The signal-based mapper Uncalled [13] follows a signal-processing, seed-mapping, and seed-clustering paradigm. The *k*-mer boundaries are inferred from the raw signals by using an event detection algorithm [27]. The input signals are normalized such that their mean and standard deviation over a rolling window match the ONT pore model [28]. The value of an event is then the mean of the signal values it covers. Next, these events are mapped to all possible *k*-mers, and the locations of those *k*-mers are identified in the reference using FM index [7]. It is ensured that the adjacent *k*-mers overlap by *k* − 1 bases. Hence, the search space forms a forest, where the positions of the signal in the reference are represented as a root-to-leaf path.

The other class of methods involves directly mapping the signals without converting them into nucleotides. Instead, the reference sequence is converted to a sequence of expected signal values using an input pore model. Furthermore, the signal values are quantized, and consecutive values are hashed [9], or organized in a k-d tree [35] and used as seeds for read mapping. Sigmoni [24] uses matching statistics over the signal values to do read classification. Working directly with noisy signals is challenging. For example, if provided with only a single chunk of signal as input, RawHash2 [9] and Sigmap [35] are unable to classify 95% and 86% of reads, respectively [24]. Hence, avoiding basecalling can prove to be more costly.

In this work, we propose two strategies to improve both classification time and F1 score compared to the existing techniques that process raw signals directly. We implement both strategies on top of an existing signal-based read mapper RawHash2 [9]. First, we use only the user-specified target regions as the reference for mapping all reads instead of the complete reference genome. This significantly improves the mapping time. The mapping outcome is used to infer the classification label. Second, we leverage alignment statistics computed by the read mapper to improve the F1 score. This is done by training an XGBoost classifier [5] over the features available from the mapped data.

We implement these methods in our tool, NanoLabel (https://github.com/at-cg/NanoLabel). We use NanoLabel to classify reads from publicly available ONT R10 human datasets. We select a panel of 258 hereditary cancer genes [19] as our target. The experimental evaluations show 80*×* improvement in the classification time, and an improvement by 0.10–0.25 units in the F1 score over RawHash2 [9] for classifying signal data using the first five chunks. In addition, we conduct an experiment on a simulated dataset with controlled noise, which provides insights into potential avenues for further improvement.

## 2. Preliminaries

### 2.1 RawHash2

RawHash2 is a state-of-the-art tool for signal-to-reference mapping [8, 9]. It is the signal-domain analogue of Minimap2 [15]. Although nucleotides can be represented by only four characters, the signal values span a much wider range. To bridge this gap, RawHash2 processes both the reference sequence and the signal output from a nanopore sequencing device to bring the two representations closer together. The reference sequence is converted to an ideal signal representation using an input pore model. Next, the signals (both from the reference and the input) are first normalized and then converted to a compact representation. For more details on the indexing procedure, see Sections 2.2-2.4 in [8].

RawHash2 then uses the seed hits between the reference genome and the query signal (known as anchors) to find the candidate mapping positions on the reference by a process called chaining [1]. Subsets of anchors (chains) that satisfy some optimisation criteria are computed using dynamic programming. The optimal chaining score is also used to compute the quality value (confidence score) of the mapping. RawHash2 provides a feature to evaluate the regions between the consecutive anchors using dynamic time warping. This helps in finding better candidate positions for mapping at the cost of higher runtime. The user can configure a maximum prefix length of the input signal using the `--max-chunks` flag that the algorithm is allowed to use to determine the best signal mapping.

### 2.2. Extreme Gradient Boosting (XGBoost) classifier

XGBoost [5] is a fast, scalable implementation of gradient-boosted decision trees. The model builds trees sequentially, where the current tree is trained on the residuals of the previous iterations. Split evaluation and node construction within each tree is heavily parallelized for speed. It often outperforms deep learning models over tabular datasets and requires much less tuning [25].

## 3. Methods

### 3.1. Overview

We propose NanoLabel, a fast and accurate real-time classifier for nanopore signals. NanoLabel uses RawHash2 to determine whether the input signal comes from the target versus non-target regions. Raw signals from ONT sequencing devices are noisy and hence challenging to process. Consequently, RawHash2 achieves a mapping F1 score of just 0.76 on the human genome while requiring high runtime for full signal mapping [9]. This makes it infeasible to use RawHash2 directly for adaptive sampling on human data. An ideal algorithm should accurately and quickly make decisions using fewer chunks.

We make significant progress in this direction compared to previous works. We first identify that using only the target regions as the reference significantly improves mapping time. But the use of an incomplete reference for mapping increases the number of false positives, hence reducing the overall classification F1 score. Next, we observe that the signal mapping statistics available in the output (PAF) file differ significantly between the false-positive and true-positive data. Hence, for the given target regions, these statistics can be exploited to train a supervised classification model to reduce the false positive rate.

To support adaptive sampling in a streaming setting, we train five separate classification models corresponding to signal lengths ranging from one to five chunks. This is a deliberate design choice in NanoLabel, made under the hypothesis that a fine-grained, signal-length-specific training model improves classification performance. Users can select an appropriate model based on how many chunks they wish to use for classification. Figure 1 shows an overview of our method when using x chunk as the input signal length. As pore models improve in the future, the signal mapping algorithm, RawHash2, can be seamlessly replaced with any other signal-mapping tool in our workflow. The following subsections describe our method in detail.

**Fig. 1.**
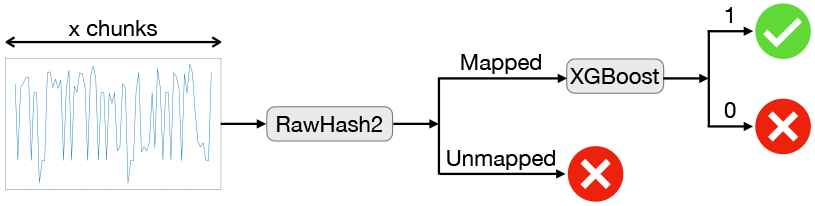
Flowchart showing the overview of NanoLabel. Assuming that the length of the input signal is x chunks (x ∈ {1, 2, 3, 4, 5}), a read is predicted as on-target (depicted by a tick) if it is mapped and the classification label predicted by XGBoost is 1. A read is predicted as off-target (depicted by a cross) if it is either unmapped or the classification label predicted for it is 0.

### 3.2. Using target regions as reference

The human genome, being highly repetitive, poses a challenge for any signal mapping tool. This may increase the mapping time and the number of candidate locations to which a signal maps. This problem can be mitigated by considering only the target regions (e.g., a set of user-specified genes) as the reference.

RawHash2 is designed for signal mapping, that is, finding positions where the signal maps on the reference genome, whereas adaptive sampling can be achieved with a binary classifier. Classification can be performed by using RawHash2 with only the target regions as the reference. This idea has been used in [13] for R9 data to enrich a 148-gene hereditary cancer panel by rejecting all the reads that did not map within the first 3 seconds. The authors claim to achieve 3.6*×* and 5.5*×* enrichment for libraries prepared with and without shearing, respectively. This is compared with a control experiment, which was also run using the same gene reference. We conduct a more rigorous analysis in this setting using R10 data and demonstrate its potential to make adaptive sampling scalable through signal-based mappers. ONT phased out the older R9 flow cells in favor of the current R10 flow cells in 2022. The signal characteristics of R10 flow cells differ substantially from those of R9 [34].

We next define the evaluation metrics with respect to our restricted reference comprising only the target regions. It is useful to have these definitions before discussing further details of the method. The reads belonging to the target region and mapped are marked as *True Positives* (abbreviated as TP), and the reads belonging to the non-target region and not mapped are marked as *True Negatives* (abbreviated as TN). Consequently, *False Positives* (FP) will be the reads not belonging to the target region and mapped. Therefore, |*FP* | equals the total count of reads not belonging to the target minus |*TN* |. Similarly, *False Negatives* (FN) will be the reads belonging to the target and not mapped. |*FN* | equals the total count of reads belonging to the target minus |*TP* |. Therefore, precision (P), recall (R), and F1 score can be written as follows:

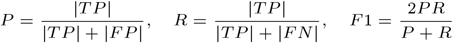

Ideally, if a read belongs to the target, it should be mapped to some position on the target. And, if it does not belong to the target, it should remain unmapped. A smaller search space in the restricted reference should ideally yield fewer spurious anchors, thereby increasing TP. This should also reduce mapping time by reducing the number of anchors and the dynamic-programming runtime during the extension step. In practice, our observations are consistent with this expectation (also shown later in Results). The fraction of reads that are correctly marked as on-target increases. However, the fraction of reads that are incorrectly marked as on-target (i.e., FP) also increases. This behavior stems from the underlying mapping algorithm, which assumes a complete reference and that every read originates from some locus within it. An off-target read may still be mapped to the locus with the highest alignment or chaining score. Due to the strong class imbalance between on-target and off-target reads, even a small increase in false positives can substantially degrade the overall F1 score. We next demonstrate how false mappings can be effectively reduced with negligible computational overhead, which is a key novelty of this work.

### 3.3. Reducing false positive mappings

For a given set of input signals, all mapping tools, including RawHash2, report their results in a standard output format (PAF). The PAF file contains key mapping parameters for each mapped read, such as the quality score, mapping time, and the number of chunks used. We further modified the RawHash2 source code to emit additional features, such as alignment scores, in the output.

When we use only the target regions as the reference, we observe that the distributions of these parameters differ significantly between reads that truly originate from the target regions and off-target reads that are nevertheless mapped (see Figure 2). We exploit this fact by training an XGBoost model on these parameters for *only* the mapped data (ref. Supplementary Section A for the set of hyperparameters used for training). We generated the ground-truth classification labels (on-target, off-target) by using Minimap2 on basecalled data. We ensured that the training data has an equal number of positive and negative samples. The data is then randomly shuffled and split into training and validation sets with a 7:3 ratio. We gathered a total of 19 features associated with mapping statistics that help improve the final F1 score (ref. Supplementary Section B for the complete list of features). We chose XGBoost because it outperformed other models, such as LightGBM, Extra Trees, and Random Forest, in our preliminary experiments. During inference, if RawHash2 reports multiple mappings for a read, only the mapping with the highest mapping quality is passed to the XGBoost model.

**Fig. 2.**
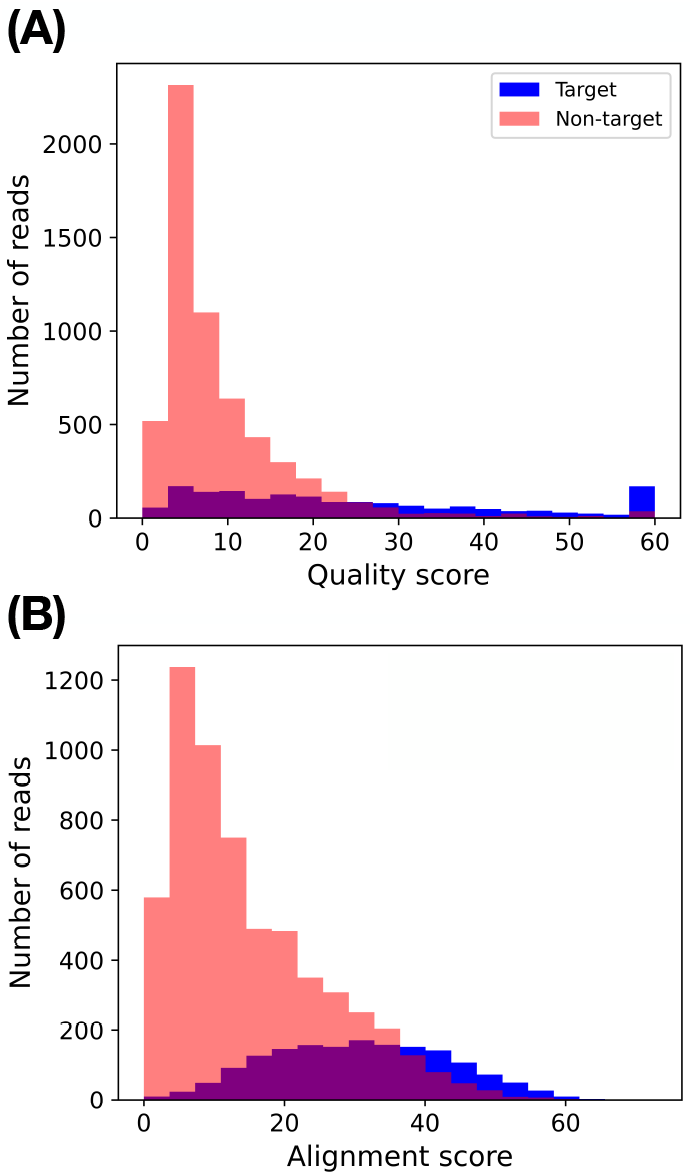
Histogram showing the distribution of (A) quality score (i.e., confidence score) and (B) alignment score for mapped reads originating from target regions (shown in blue) and non-target regions (shown in red). In this experiment, only the first signal chunks from dataset D1 are used for mapping (Table 1).

For a given target panel, both training data generation and model training are performed as a preprocessing step. This preprocessing can be done before using ONT’s adaptive sampling functionality. Overall, this strategy is agnostic to the underlying signal mapping algorithm. Hence, it can be integrated with other mapping tools in the future.

**Table 1.** Statistics of the real and simulated datasets used in our benchmark.

| Dataset (URL) | Genome | Coverage | N50 read length | # Reads |  |  |  |
| --- | --- | --- | --- | --- | --- | --- | --- |
|  |  |  |  | Input | After processing | # Target | # Non-target |
| D1 ([31]) | HG001 | 14.62× | 28,366 | 2,385,918 | 2,295,325 | 17,542 | 2,277,783 |
| D2 ([31]) | HG001 | 17.53× | 28,305 | 2,834,704 | 2,719,122 | 20,707 | 2,698,415 |
| D3 ([32]) | HG002 | 49.30× | 29,360 | 7,463,014 | 7,017,825 | 52,536 | 6,965,289 |
| D4 ([33]) | HG005 | 27.77× | 23,956 | 4,137,223 | 3,869,800 | 30,171 | 3,839,629 |
| D5 (Simulated) | GRCh38 | 9.99× | 20,052 | 2,042,623 | 2,042,623 | 15,990 | 2,026,633 |
| D6 (Simulated) | GRCh38 | 10.00× | 20,081 | 2,042,807 | 2,042,807 | 16,126 | 2,026,681 |

### 3.4. Data augmentation

The target regions would usually be much smaller in length than the full reference, which means gathering sufficient training data can be challenging. First, there is a severe class imbalance (Results section, Table 1). Second, RawHash2 has a low mapping rate when only a small number of chunks are available as input (Results), whereas we require mapped data for training. As a result, the number of reads mapped may be insufficient to construct a balanced training dataset. One easy way to address this demand would be to use simulated data [10] for training. However, we observed in our preliminary experiments that there is a mismatch between the noise characteristics of simulated and real nanopore signals. To address this limitation, we propose a novel data augmentation strategy (Algorithm 1) that enables the generation of sufficient training data from real sequencing signals.

For the 258-cancer-gene panel used in our study, the count of non-target reads is more than 100 times the count of reads coming from the target region in all the datasets (Table 1). Therefore, we only augment the count of reads coming from the target region. The augmentation procedure is tailored to the signal length used by NanoLabel for classification, which ranges from one to five chunks across the five classification models. Suppose we want to augment when the input signal has *c* chunks, and each chunk has *s* current measurements.

#### Algorithm 1

Data augmentation procedure and training of XGBoost models

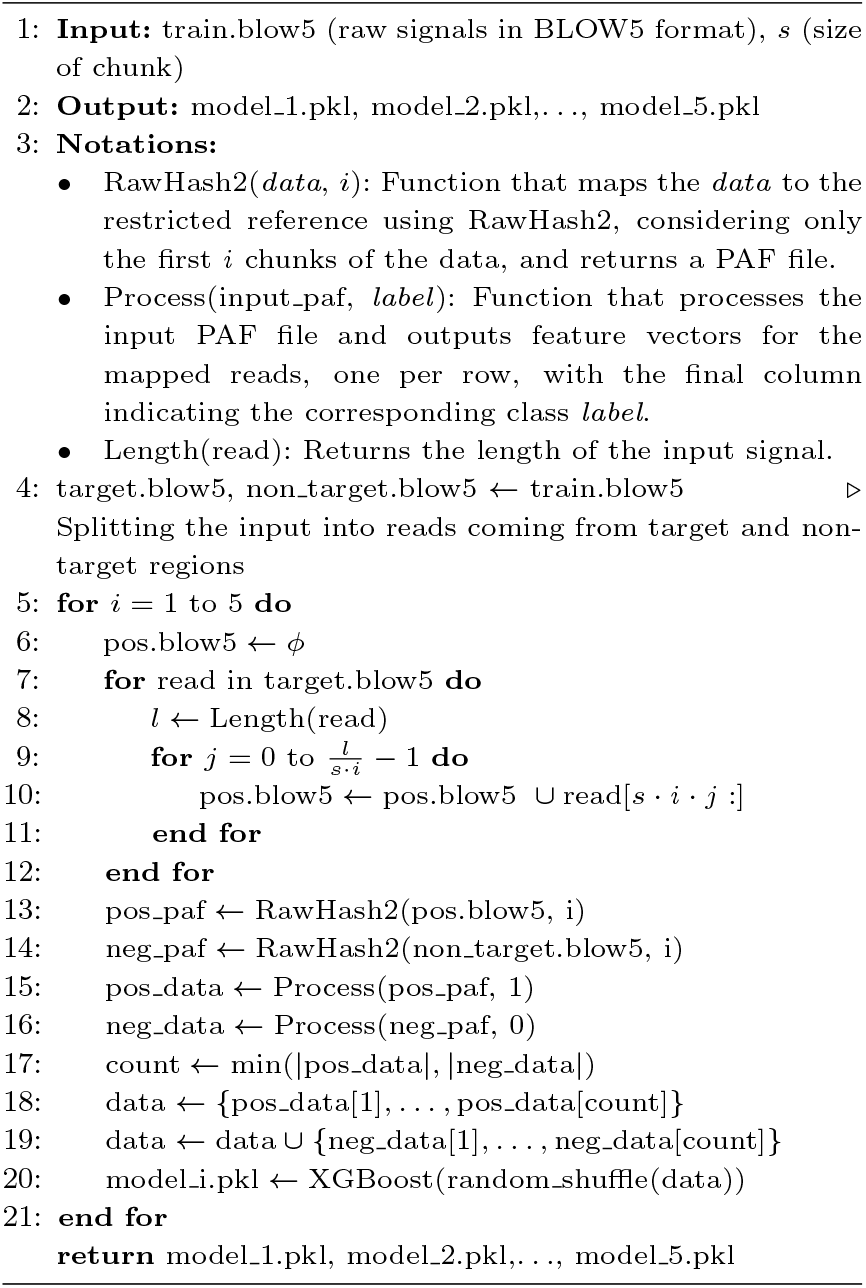

We augment the signal associated with a single read by taking suffixes starting after every *s · c* values (Line 10 in Algorithm 1). The intuition is that RawHash2 will use only the initial *s · c* values for mapping when we configure its `--max-chunks` parameter setting to *c*. Therefore, full signals that belong to the target region can not be exploited. By performing the proposed augmentation, we utilize the input data to its maximum potential and avoid redundancy in the output, which can hinder the training process. The proposed augmentation strategy is highly effective when the target regions are much smaller and sufficient training data is hard to gather.

## 4. Experiments

All our experiments are done on a dual-socket Intel Xeon Gold 6248R server with 2*×*24 cores and 750 GB RAM.

### Evaluated methods

We set up experiments to compare three methods: RawHash2 using the complete reference (RH2-CR), RawHash2 using a restricted reference (RH2-RR), and the method proposed in this work, NanoLabel. We do not include other signal-level classification tools, such as Uncalled [13], Sigmap [35], and Sigmoni [24] in our comparison because these tools are either not designed^∗^ or tuned^†^ for the R10 data.

For all evaluated methods (RH2-CR, RH2-RR, and NanoLabel), reads that do not map to the complete reference (in RH2-CR) or to the restricted reference (in RH2-RR and NanoLabel) are assumed to originate from non-target regions. This choice is made due to the strong class imbalance in all datasets, where a read is more likely to belong to the negative class. Treating unmapped reads as on-target in either method would reduce their classification F1 score.

In RH2-CR, mapped reads are classified as on-target versus off-target based on their mapping positions in the complete reference genome. In RH2-RR, all mapped reads are classified as on-target. NanoLabel performs the classification of a mapped read using the XGBoost inference (ref. Section 3.3).

### Datasets

As target regions in the human genome, we use a panel of 258 genes from the Oxford Nanopore Hereditary Cancer Panel [19]. To construct a restricted reference, we extracted the 258 gene sequences from the GRCh38 reference genome by using gene annotations from Gencode (release 48). We also included 10 kbp flanking sequence upstream and downstream of each gene (20 kbp of flanking sequence in total per gene). The flanking sequence is included to cover those reads that start outside but could extend into a gene.

We evaluate all methods using publicly available whole-genome sequencing datasets from the HG001, HG002, and HG005 human genomes (Table 1). These datasets were sequenced using PromethION R10.4.1 flow cells [30]. The HG001, HG002, and HG005 genomes represent individuals from different ancestral backgrounds, allowing us to assess the robustness of our methods across diverse genetic variation. We split the HG001 dataset further into two disjoint subsets, with one reserved for training. We also generate simulated data from the GRCh38 reference genome using Squigulator [10]. We utilize different subsets of data for training and testing to demonstrate the generalizability of our method (see Table 2).

**Table 2.** Combinations of the training and testing datasets used in the experiments.

| Experiment number | Training data | Testing data |
| --- | --- | --- |
| 1 | D1 | D2, D3, D4 |
| 2 | D1 + D2 | D3, D4 |
| 3 | D5 | D6 |

### Labelling the training and test datasets

The real sequencing datasets are available in POD5 format. We convert these files to BLOW5 format using Blue-crab [11], as BLOW5 format facilitates efficient file manipulation and collection of signal-level statistics [22]. The original POD5 files are basecalled using Dorado’s *super-accurate* (SUP) model [29].

Reads shorter than 1 kbp are removed from both the BLOW5 files and the corresponding FASTA/FASTQ files, as short reads are unlikely to affect target enrichment and are therefore excluded from the classification analysis. To generate ground-truth labels, reads in the FASTQ files are aligned to the GRCh38 reference genome using Minimap2 [16]. We use the `--secondary=no` option in Minimap2 to suppress secondary alignments. Unmapped reads and the reads having supplementary alignments are excluded. Counts of the remaining reads after processing are shown in Table 1. A read is labeled as positive if and only if its complete alignment lies within one of the target intervals, where each target interval includes the gene together with its flanking regions. Ground-truth labels from the training data are used for supervised learning in NanoLabel. Labels from the test data are used exclusively to evaluate the accuracy of the methods.

### Training data augmentation

The training datasets are further augmented to obtain sufficient, class-balanced data for training the XGBoost models (ref. Section 3.4). The total size of data after augmentation is provided in Table 4. We observe approximately 3*×* growth in the training data after applying our augmentation strategy (Tables 3, 4). For shorter signals (e.g., a single chunk), the training set contains fewer reads compared to longer signals. This is due to RawHash’s low mapping rate when only one or two chunks are provided as input. We only include mapped reads in training data.

**Table 3.** The size of the training datasets in Experiment 1 without performing any augmentation. Each dataset has an equal count of positively and negatively labelled data.

| Signal length<br>(number of chunks) | 1 | 2 | 3 | 4 | 5 |
| --- | --- | --- | --- | --- | --- |
| # Reads | 3,244 | 11,612 | 18,802 | 23,280 | 26,218 |

**Table 4.** The size of the training datasets across the three experiments after augmentation. Each dataset has an equal count of positively and negatively labelled data.

| Signal length<br>(number of chunks) | Exp. 1 | Exp. 2 | Exp. 3 |
| --- | --- | --- | --- |
| 1 | 11,938 | 25,486 | 66,032 |
| 2 | 30,958 | 66,642 | 133,656 |
| 3 | 50,692 | 110,472 | 185,868 |
| 4 | 69,448 | 151,088 | 228,510 |
| 5 | 87,298 | 190,288 | 200,386 |

### RawHash2 parameters

We use the default parameter settings for RawHash2, which are recommended in its documentation for large reference genomes. To further tailor RawHash2 to the restricted reference, we explored a range of parameter configurations and found that modifying most parameters led to substantial degradation in either mapping runtime or accuracy. The only exception is enabling the dynamic time warping (DTW)-based extension, which we activate for the restricted reference to further improve mapping accuracy. RawHash2 is allocated 48 threads for faster execution.

RawHash2 additionally requires a pore model that defines the correspondence between signal events and *k*-mers. For real datasets, we use a custom pore model generated with Uncalled4 [14]. For simulated data, we use the default pore model provided by ONT, as the simulated data was generated using the same model. The RawHash2 commands used for reference index construction and signal mapping are provided in Supplementary Section F.

### Evaluation metrics

For evaluation on the test datasets, we consider read prefixes of varying lengths, ranging from one to five chunks, with each chunk comprising approximately 4000 current measurements. In other words, we measure the classification accuracy and runtime of the methods under varying signal lengths. Using more than five chunks would undermine the objective of adaptive sampling, as a substantial portion of the DNA molecule would have already translocated through the pore.

We evaluate classification performance using four metrics: the F1 score, the ratio of true positives to total positives (FTP), the ratio of true negatives to total negatives (FTN), and the classification time. The performance of different methods is easier to interpret using the FTP and FTN metrics; therefore, we choose these measures over precision and recall.

### Results

We first show the F1 score achieved by the XGBoost models on the validation split of the training data (which only includes mapped data) in Table 5. All values are more than 0.9. Hence, the model learns the mapping features of the unseen dataset well. Next, we discuss the performance on test datasets.

**Table 5.** F1 scores achieved by the XGBoost models on the validation split. The validation data is 30% of the total input data (Table 4).

| Signal length<br>(number of chunks) | Exp. 1 | Exp. 2 | Exp. 3 |
| --- | --- | --- | --- |
| 1 | 0.93 | 0.93 | 0.93 |
| 2 | 0.93 | 0.93 | 0.95 |
| 3 | 0.92 | 0.93 | 0.95 |
| 4 | 0.93 | 0.93 | 0.96 |
| 5 | 0.93 | 0.93 | 0.96 |

#### *Experiment* 1

In Experiment 1, we evaluate the methods on publicly available whole-genome sequencing datasets from the HG001 (D2), HG002 (D3), and HG005 (D4) human genomes.

The corresponding results are reported in Tables 6, 7, and 8 respectively. Evaluating performance across three distinct genomes provides a robust assessment of method generalizability.

**Table 6.** Benchmarking results on dataset D2 (HG001 sequencing data) in Experiment 1. Columns labeled FTP and FTN denote the ratios of true positives to total positives and true negatives to total negatives, respectively. The best F1 scores are highlighted in bold.

| Signal length<br>(number of chunks) | RH2-CR |  |  | RH2-RR |  |  | NanoLabel |  |  |
| --- | --- | --- | --- | --- | --- | --- | --- | --- | --- |
|  | F1 | FTP | FTN | F1 | FTP | FTN | F1 | FTP | FTN |
| 1 | 0.02 | 0.010 | 0.999 | 0.12 | 0.086 | 0.997 | **0.13** | 0.070 | 0.999 |
| 2 | 0.15 | 0.083 | 0.999 | 0.29 | 0.318 | 0.993 | **0.41** | 0.279 | 0.999 |
| 3 | 0.33 | 0.203 | 0.999 | 0.35 | 0.524 | 0.988 | **0.59** | 0.473 | 0.999 |
| 4 | 0.48 | 0.337 | 0.999 | 0.36 | 0.659 | 0.984 | **0.68** | 0.600 | 0.998 |
| 5 | 0.58 | 0.452 | 0.999 | 0.35 | 0.742 | 0.980 | **0.72** | 0.683 | 0.998 |

**Table 7.** Benchmarking results on dataset D3 (HG002 sequencing data) in Experiment 1. Columns labeled FTP and FTN denote the ratios of true positives to total positives and true negatives to total negatives, respectively. The best F1 scores are highlighted in bold.

| Signal length<br>(number of chunks) | RH2-CR |  |  | RH2-RR |  |  | NanoLabel |  |  |
| --- | --- | --- | --- | --- | --- | --- | --- | --- | --- |
|  | F1 | FTP | FTN | F1 | FTP | FTN | F1 | FTP | FTN |
| 1 | 0.03 | 0.013 | 0.999 | 0.13 | 0.088 | 0.997 | **0.14** | 0.074 | 0.999 |
| 2 | 0.15 | 0.085 | 0.999 | 0.30 | 0.313 | 0.993 | **0.41** | 0.278 | 0.999 |
| 3 | 0.32 | 0.201 | 0.999 | 0.36 | 0.511 | 0.989 | **0.59** | 0.464 | 0.999 |
| 4 | 0.46 | 0.323 | 0.999 | 0.36 | 0.639 | 0.985 | **0.67** | 0.584 | 0.998 |
| 5 | 0.57 | 0.434 | 0.999 | 0.35 | 0.720 | 0.981 | **0.71** | 0.662 | 0.998 |

**Table 8.** Benchmarking results on dataset D4 (HG005 sequencing data) in Experiment 1. Columns labeled FTP and FTN denote the ratios of true positives to total positives and true negatives to total negatives, respectively. The best F1 scores are highlighted in bold.

| Signal length<br>(number of chunks) | RH2-CR |  |  | RH2-RR |  |  | NanoLabel |  |  |
| --- | --- | --- | --- | --- | --- | --- | --- | --- | --- |
|  | F1 | FTP | FTN | F1 | FTP | FTN | F1 | FTP | FTN |
| 1 | 0.03 | 0.013 | 0.999 | 0.13 | 0.091 | 0.997 | **0.14** | 0.077 | 0.999 |
| 2 | 0.16 | 0.089 | 0.999 | 0.31 | 0.325 | 0.993 | **0.43** | 0.290 | 0.999 |
| 3 | 0.34 | 0.211 | 0.999 | 0.37 | 0.537 | 0.989 | **0.61** | 0.494 | 0.999 |
| 4 | 0.49 | 0.344 | 0.999 | 0.38 | 0.669 | 0.985 | **0.69** | 0.616 | 0.998 |
| 5 | 0.60 | 0.463 | 0.999 | 0.37 | 0.753 | 0.981 | **0.74** | 0.698 | 0.998 |

When using RawHash2 with the complete reference (RH2-CR), the FTP remains below 0.1 for two chunks and increases to approximately 0.5 with five chunks, limiting the F1 score to around 0.6 even at five chunks. The low F1 scores at shorter prefixes arise because RawHash2 fails to map most reads to the complete reference when only a few chunks are available. This behavior is consistent with prior observations [24]. Using a restricted reference (RH2-RR) improves FTP by at least 60%, validation split. The validation data is 30% of the total input data (Table 4). but this comes at the cost of reduced FTN. As discussed earlier, this trade-off arises from the inherent behavior of RawHash2’s read-mapping algorithm, which attempts to assign each read to the highest-scoring match in the reference. Overall, this reduces the F1 score relative to RH2-CR, with performance saturating beyond three chunks.

Incorporating the XGBoost model in NanoLabel improves the classification of mapped reads, leading to marked gains in F1 score. Notably, NanoLabel achieves a higher F1 score with three chunks than RH2-CR does with five chunks, demonstrating strong generalization to unseen data.

We also measure classification time, which would be the time required to decide whether a read should be accepted or rejected. By refining RawHash2’s decisions through learned classification of mapped reads, NanoLabel not only improves accuracy but also significantly reduces classification time compared to RH2-CR (Table 9). The total classification time of NanoLabel includes mapping of reads to the restricted reference and XGBoost inference, which is effectively identical to that of RH2-RR up to two decimal places. This implies that the time used for XGBoost inference is negligible. In contrast, RH2-CR requires at least 80*×* more time than either RH2-RR or NanoLabel.

**Table 9.** Classification time of the methods (in ms) on the test datasets in Experiment 1. The best values are highlighted in bold.

| Signal<br>(#chunks) | HG001 |  |  | HG002 |  |  | HG005 |  |  |
| --- | --- | --- | --- | --- | --- | --- | --- | --- | --- |
|  | RH2-CR | RH2-RR | NanoLabel | RH2-CR | RH2-RR | NanoLabel | RH2-CR | RH2-RR | NanoLabel |
| 1 | 461.6 | **4.8** | **4.8** | 491.1 | **4.9** | **4.9** | 464.8 | **4.8** | **4.8** |
| 2 | 1179.5 | **12.8** | **12.8** | 1295.1 | **12.0** | **12.0** | 1269.7 | **11.8** | **11.8** |
| 3 | 2062.2 | **23.7** | **23.7** | 2297.5 | **21.6** | **21.6** | 2351.4 | **21.6** | **21.6** |
| 4 | 3370.2 | **36.4** | **36.4** | 3412.1 | **34.4** | **34.4** | 3379.6 | **34.6** | **34.6** |
| 5 | 4309.6 | **52.7** | **52.7** | 4646.6 | **49.7** | **49.7** | 4435.5 | **49.4** | **49.4** |

**Table 10.** Benchmarking results on simulated dataset D6 in Experiment 3. Columns labeled FTP and FTN denote the ratios of true positives to total positives and true negatives to total negatives, respectively. The best F1 scores are highlighted in bold.

| Signal length<br>(number of chunks) | RH2-CR |  |  | RH2-RR |  |  | NanoLabel |  |  |
| --- | --- | --- | --- | --- | --- | --- | --- | --- | --- |
|  | F1 | FTP | FTN | F1 | FTP | FTN | F1 | FTP | FTN |
| 1 | 0.58 | 0.442 | 0.999 | 0.36 | 0.677 | 0.983 | **0.70** | 0.628 | 0.998 |
| 2 | **0.80** | 0.744 | 0.999 | 0.29 | 0.870 | 0.966 | 0.79 | 0.832 | 0.997 |
| 3 | **0.86** | 0.855 | 0.998 | 0.24 | 0.926 | 0.953 | 0.80 | 0.886 | 0.997 |
| 4 | **0.88** | 0.903 | 0.998 | 0.21 | 0.950 | 0.943 | 0.80 | 0.911 | 0.997 |
| 5 | **0.90** | 0.928 | 0.998 | 0.19 | 0.963 | 0.934 | 0.79 | 0.915 | 0.996 |

For five chunks, NanoLabel completes classification in approximately 50 ms, which is an order of magnitude faster than the time required for a pore to read one chunk. Performing fast and accurate classification can positively impact the enrichment of target molecules during adaptive sampling.

#### *Experiment* 2

In Experiment 2, we investigate whether increasing the training data improves NanoLabel’s performance during inference on the test data. The results in Supplementary Tables S2 and S3 show that the training dataset used for Experiment 1 is already sufficient to capture nearly all the diversity in the feature vectors generated by RawHash2 over the chosen target regions. That is why, even after doubling the training data (Table 4), the accuracy metrics remain almost the same. We conclude that a dataset of the order of 10^4^ samples is sufficient to train the XGBoost model.

#### *Experiment* 3

In Experiment 3, we test our method using simulated data. The simulated signals are generated using Squigulator, which relies on traditional pore models and Gaussian noise [10]. Due to the simplicity of these models, the simulated data do not fully capture the complexity of real nanopore signals. Nevertheless, this experiment is interesting because the classification methods perform differently on simulated data.

RH2-CR performs significantly better on simulated data, with F1 and FTP values reaching 0.9 for 5 chunks. This is because RawHash2 is able to map a larger number of reads across all signal lengths. This also suggests that handling noise in real data using computationally lightweight signal processing techniques requires further attention. When using the RH2-RR method, the mapping rate also increases. However, this also reduces the FTN values. Recall that the size of the negative class is more than 100*×* that of the positive class (Table 1). Hence, RH2-RR performs better at lower chunk counts, but its performance degrades relative to its behavior on real data as the chunk count increases.

Finally, our post-processing scheme implemented in NanoLabel performs well even with a single chunk, achieving an F1 score of 0.7. The F1 score saturates at approximately 0.8 after two chunks. This indicates a limitation of our approach, i.e., we do not correctly classify a subset of false-positive mappings. Finally, classification times on simulated data follow trends similar to those observed on real datasets: NanoLabel and RH2-RR are significantly faster than RH2-CR due to their use of a restricted reference (Supplementary Table S4).

## 5. Conclusions

In this work, we investigate adaptive sampling in a basecalling-free setting, with a focus on scenarios that have limited computational resources. Existing signal-based mappers struggle to operate reliably on noisy signals in real-time settings. We propose NanoLabel, a fast and accurate real-time signal classifier. We use RawHash2 over the target regions as the reference. While this significantly improves classification speed, it also introduces false positives. Due to the strong class imbalance toward the negative class, the overall F1 score degrades. We mitigate this problem by distinguishing false positives from true positives using an XGBoost classifier trained on mapping-derived features.

We evaluate NanoLabel to classify reads sampled from the 258-gene hereditary cancer panel [19] versus the remaining human genome. We observe significant improvements in both accuracy and runtime compared to directly using RawHash2 with the complete reference (RH2-CR). For example, on real sequencing datasets, NanoLabel achieves an F1 score of 0.6 using 3 chunks of data. This performance is achieved by RH2-CR using 5 chunks. Moreover, NanoLabel reduces classification time by nearly two orders of magnitude relative to RawHash2 on the complete reference. Overall, these results demonstrate the potential utility of the proposed approach for adaptive sampling and target enrichment on large genomes.

We identify a few promising directions for future work. First, it may be useful to explore and integrate more features for distinguishing between off-target and on-target mapped reads, e.g., matching statistics [24]. Second, developing alternative signal-level mapping algorithms that are better suited for references restricted to target regions may reduce reliance on supervised classification. Third, designing more accurate, lightweight statistical or machine learning models for raw signal processing tasks, such as detecting *k*-mer event boundaries in the raw signal, would improve the effectiveness of downstream mapping and classification tasks.

## Acknowledgments

We thank Ishaan Gupta, Chandra Sekhar Seelamantula, and Prathosh A.P. for useful discussions. We also thank Can Firtina and Vikram Shivakumar for responding to our queries on GitHub. This research is supported in part by funding from the DBT/Wellcome Trust India Alliance (IA/I/23/2/506979). We used computing resources provided by the National Energy Research Scientific Computing Center (NERSC), USA.

## Footnotes

∗ https://github.com/skovaka/UNCALLED/issues/36#issuecomment-909321265

† https://github.com/vikshiv/sigmoni/issues/5

