## Supplementary file for "NanoLabel: A fast and accurate real-time nanopore signal classifier"

### Supplementary Document

Comments are indicated using #*.

#### A. Hyperparameters used for training the XGBoost model

```
clf = XGBClassifier(random_state = 0, n_estimators = 10)
gbm_param_grid = {'eta' : [0.01, 0.05, 0.1], 'max_depth' : [3, 5, 7, 9]}
clf = GridSearchCV(estimator = clf, param_grid = gbm_param_grid, scoring = 'f1', cv = 5, verbose = 1)
```

#### B. Features used for training the XGBoost model

```
query_aligned_length, reference_aligned_length, fraction_query_aligned_length,
fraction_reference_aligned_length, quality_score, mapping_time, number_of_chunks_used,
total_query_length, number_of_mappings.
```

Also, for the first two mappings of a query signal, the following parameters:

```
number_of_minimizers, dynamic_programming_alignment_score, best_alternate_mapping_score,
number_of_suboptimal_mappings, alignment_score
```

#### C. Command for getting the location of all genes

```
## Downloading the complete file from the Gencode website:

wget "https://ftp.ebi.ac.uk/pub/databases/gencode/Gencode_human/release_48/gencode.v48.annotation.gtf
    .gz"


## Extracting the positions of all genes:

zcat gencode.v48.annotation.gtf.gz |
awk '$3=="gene" {
    match($0, /gene_name "([^"]+)"/, arr);
    if (arr[1] != "") {
        OFS=" ";
        print $1, $4-1, $5, arr[1];
    }
}' > all_genes.bed
```

#### D. Command for generating simulated data

```
squigulator hg38_ref.fa -x dna-r10-prom -r 15000 -f 10 -t 48 --sample-rate 4000 -o $SQUIGGLE_OUT -q
    $FASTA_OUT
```

#### E. Processing POD5 files

```
## Converting POD5 to BLOW5 format:

blue-crab p2s $POD5_PATH -d $BLOW5_PATH
slow5tools merge $BLOW5_PATH/*.blow5 --to blow5 -t 48 -o $BLOW5_PATH/reads.blow5
```

```
## Basecalling the POD5 file:

dorado basecaller sup $POD5_PATH --emit-fastq > $FASTQ_PATH


## Mapping the FASTQ file:

minimap2 -cx map-ont -t 48 --secondary=no hg38_ref.fa $FASTQ_PATH > $PAF_PATH


## Removing reads of length less than 1000:

seqtk seq -L 1000 $FASTQ_PATH > $PROCESSED_FASTQ_PATH  ## similar command for fasta files

// custom script to remove reads of less than 1000 from reads.blow5 and $PAF_PATH using read_ids from
     $PROCESSED_FASTQ_PATH
```

#### F. RawHash2 commands

```
if [ $DATA is simulated ]; then
    KMER_MODEL=rawhash2/extern/kmer_models/dna_r10.4.1_e8.2_400bps/9mer_levels_v1.txt
else
    KMER_MODEL=rawhash2/extern/local_kmer_models/uncalled_r1041_model_only_means.txt
fi


## Complete reference:

### Indexing:

rawhash2 --r10 -d $REF_OUT -p $KMER_MODEL -t 48 hg38_ref.fa

### Mapping:

rawhash2 --r10 -t 48 -p $KMER_MODEL -o $RMAP_OUT -x fast --max-chunks $i $REF_OUT $DATA


## Restricted reference:

### Indexing:

rawhash2 --r10 -d $REF_OUT -p $KMER_MODEL -t 48 --store-sig restricted_ref.fa

### Mapping:

rawhash2 --r10 -t 48 -p $KMER_MODEL -o $RMAP_OUT -x sensitive --max-chunks $i --dtw-evaluate-chains
--dtw-border-constraint global $REF_OUT $DATA
```

#### G. Supplementary tables

**Table S1.** Versions of tools and libraries used in the experiments.

| Tool | Version | Link to source code |
|---|---|---|
| NanoLabel | 1.0 | https://github.com/at-cg/NanoLabel |
| RawHash2 | commit:8371101 | https://github.com/CMU-SAFARI/RawHash |
| Minimap2 | 2.26-r1175 | https://github.com/lh3/minimap2 |
| Dorado | 0.6.3 | https://github.com/nanoporetech/dorado |
| Squigulator | 0.3.0 | https://github.com/hasindu2008/squigulator |
| Slow5tools | 1.1.0 | https://github.com/hasindu2008/slow5tools |
| Blue-crab | 0.4.0 | https://github.com/Psy-Fer/blue-crab |
| Seqtk | 1.4-r122 | https://github.com/lh3/seqtk |
| Seqkit | v2.6.1 | https://github.com/shenwei356/seqkit |

**Table S2.** Benchmarking results on D3 dataset for Experiment 2. We only show the results of NanoLabel; the results for other methods remain the same as in Table 7 in the main text.

| **Signal length** | **NanoLabel** | | |
|---|---|---|---|
| (number of chunks) | **F1** | **FTP** | **FTN** |
| 1 | 0.13 | 0.073 | 0.999 |
| 2 | 0.41 | 0.280 | 0.999 |
| 3 | 0.59 | 0.466 | 0.999 |
| 4 | 0.67 | 0.588 | 0.998 |
| 5 | 0.71 | 0.667 | 0.998 |

**Table S3.** Benchmarking results on D4 dataset for Experiment 2. We only show the results of NanoLabel; the results for other methods remain the same as in Table 8 in the main text.

| **Signal length** | **NanoLabel** | | |
|---|---|---|---|
| (number of chunks) | **F1** | **FTP** | **FTN** |
| 1 | 0.14 | 0.076 | 0.999 |
| 2 | 0.43 | 0.294 | 0.999 |
| 3 | 0.62 | 0.496 | 0.999 |
| 4 | 0.70 | 0.621 | 0.998 |
| 5 | 0.74 | 0.703 | 0.998 |

**Table S4.** Decision time of the tools (in ms) for Experiment 3 in which the methods are evaluated on simulated data.

| **Signal length** (number of chunks) | **RH2-CR** | **RH2-RR** | **NanoLabel** |
|---|---|---|---|
| 1 | 1044.83 | **10.34** | **10.34** |
| 2 | 1933.11 | **27.44** | **27.44** |
| 3 | 2755.23 | **48.21** | **48.21** |
| 4 | 3355.25 | **77.86** | **77.86** |
| 5 | 3328.56 | **112.43** | **112.43** |
